# CrossTx: Cross-cell line Transcriptomic Signature Predictions

**DOI:** 10.1101/2023.01.09.523287

**Authors:** Panagiotis Chrysinas, Changyou Chen, Rudiyanto Gunawan

## Abstract

**Motivation:** Predicting the cell response to chemical compounds is central to drug discovery, drug repurposing, and personalized medicine. To this end, large datasets of drug response signatures have been curated, most notably the Connectivity Map (CMap) from the Library of Integrated Network-based Cellular Signatures (LINCS) project. A multitude of *in silico* approaches have also been formulated to leverage drug signature data for accelerating novel therapeutics. However, the majority of the available data are from immortalized cancer cell lines. Cancer cells display markedly different responses to compounds, not only when compared to normal cells, but also among cancer types. Strategies for predicting drug signatures in unseen cells—cell lines not in the reference datasets—are still lacking.

**Results:** In this work we developed a computational strategy, called CrossTx, for predicting drug transcriptomic signatures of an unseen target cell line using drug transcriptome data of reference cell lines and background transcriptome data of the target cells. Our strategy involves the combination of predictor and corrector steps. Briefly, the Predictor applies averaging (mean) or linear regression model to the reference dataset to generate cell line-agnostic drug signatures. The Corrector generates target-specific drug signatures by projecting cell line-agnostic signatures from the Predictor onto the transcriptomic latent space of the target cell line using Principal Component Analysis (PCA) and/or an Autoencoder (AE). We tested different combinations of Predictor-Corrector algorithms in an application to the CMap dataset to demonstrate the performance of our approach.

**Conclusion:** CrossTx is an efficacious and generalizable method for predicting drug signatures in an unseen target cell line. Among the combinations tested, we found that the best strategy is to employ Mean as the Predictor and PCA followed by AE (PCA+AE) as the Corrector. Still, the combination of Mean and PCA (without AE) is an attractive strategy because of its computationally efficiency and simplicity, while offering only slightly less accurate drug signature predictions than the best performing combination.

**Availability and implementation:** http://www.github.com/cabsel/crosstx

## 1. Introduction

A critical step in the drug discovery process is identifying compounds that elicit a desired activity on disease-modifying targets. In this regard, data-driven strategies play an important role in mining and integrating literature, knowledge base, and omics data (*e.g*., transcriptome), for prioritizing and matching drugs to molecular targets (Dudley et al., 2011; Jin & Wong, 2014; Kim et al., 2016; Louhimo et al., 2016; Qian et al., 2019). These strategies typically require an abundance of cellular signatures of drugs, preferably from the specific human tissue(s) affected by the disease. Of note is the Connectivity Map (CMap) dataset that contains 1.5 million human transcriptomic signatures from roughly 20,000 drug treatments and chemical perturbations for 71 different cell lines (Subramanian et al., 2017). Although impressive in its size, the large majority of drug signatures in the CMap dataset are from immortalized cancer cell lines. Unfortunately, cancer cells are known to exhibit abnormalities in drug responses in comparison to normal human tissues. Even among cancer cells, there exists a significant variability of molecular signatures in response to drugs (Subramanian et al., 2017). Surprisingly, predicting drug transcriptomic response in an unseen cell line has not received much attention and only a small number of computational strategies exist, despite the practical relevance of this prediction.

The drug signature prediction in this work is illustrated in **Fig. 1a**. The task of at hand is to predict transcriptomic drug signatures in an unseen target cell line of interest using drug signature data of reference cell lines and background data of the target cells. In practice, gene expression data of a target cell line are often available from past experiments or the literature and public databases (for example, GEO database (Edgar et al., 2002)). While the prediction in **Fig. 1a** is akin to data imputation problem, it has one key difference that makes the majority of existing imputation algorithms inapplicable: the data from the reference cell lines do not have an overlap in conditions (drug treatments) with the background data from the target cell line. Computational methods for imputation that have previously been developed and applied to the CMap dataset address a more common imputation problem that involves sparsely missing samples—this is illustrated in **Fig. 1a** as the 3-D matrix with missing samples. Several existing imputation strategies rely on the 3-D matrix representation of CMap drug signature data, where genes, cell lines, and drug conditions make up the three axes. The simplest approach is by averaging, either 1-D (across reference cell lines for the same drug) or 2-D (across drugs from the same cell line and across reference cell lines for the same drug) (Hodos et al., 2018). More sophisticated strategies rely on tensor decompositions of the 3-D matrix (Hodos et al., 2018; Iwata et al., 2019). Notably, the tensor decomposition method called TT-WOPT (Tensor-Train Weight OPTimization) (Iwata et al., 2019) is able to predict drug signatures for unseen cells (referred to as ‘missing cells’ in the original publication), which motivates its comparison to our proposed strategy.

**Fig. 1.**
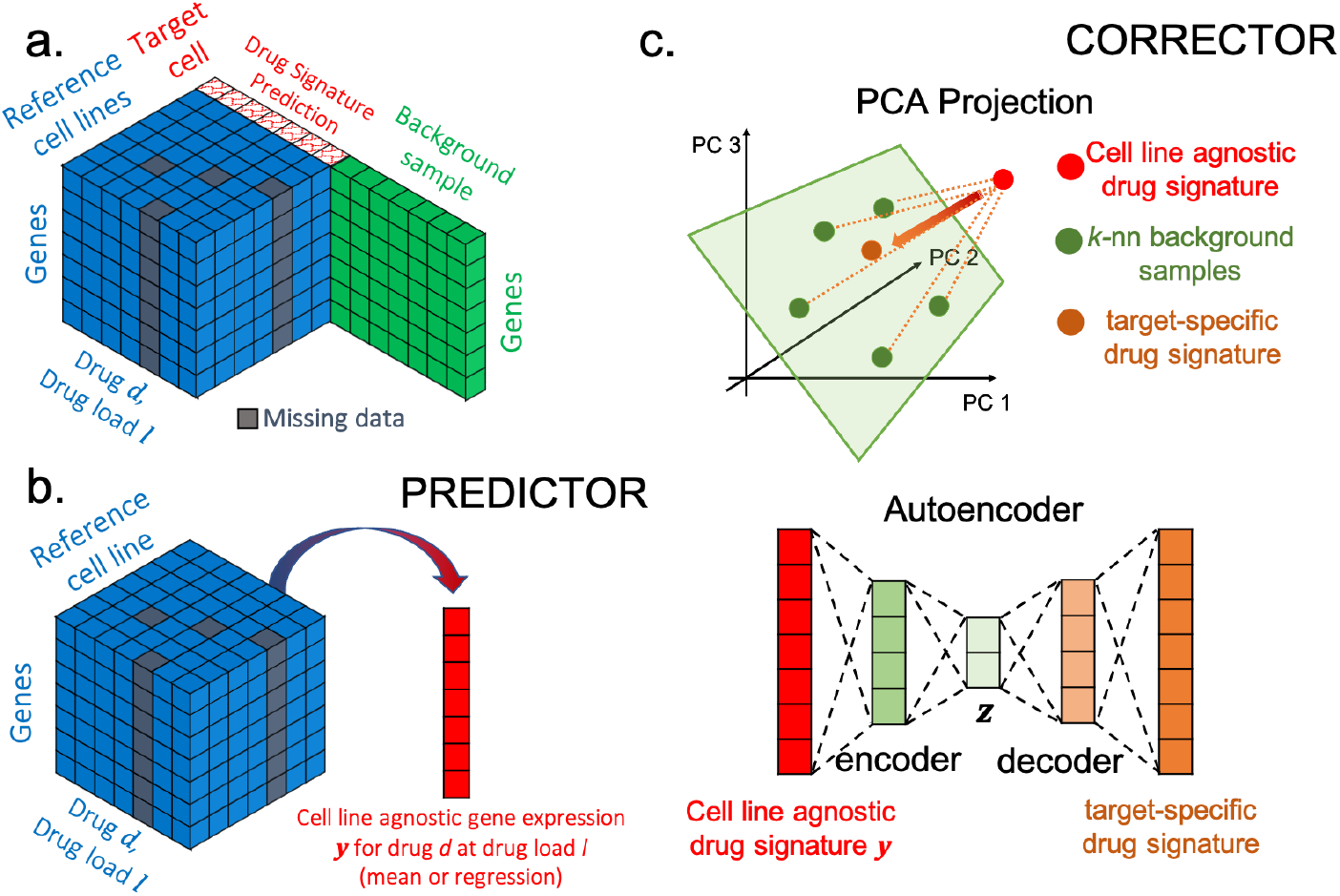
Transcriptomic Drug Signature Prediction by CrossTx. (a.) An overview of drug signature prediction in an unseen target cell line. Reference transcriptome signatures comprising gene expression data from different reference cell lines treated with various drugs and drug loads (shaded in blue) are displayed as a 3-D matrix. The combinations of drug and drug loads may not be the same across the reference cell lines, displayed as missing data samples (shaded in grey). Background gene expression data of the target cell line (shaded in green) do not have any overlapping conditions (drug treatements) with the reference signatures. Drug signature to be predicted is highlighted in red. (b.) The Predictor generates cell line-agnostic gene expression by averaging or linear regression model. (c.) The Corrector projects cell line-agnostic signatures from the Predictor using PCA and/or AE, to produce target-specific gene expression signatures.

Another group of methods are based on supervised learning strategy where a linear regression model or machine learning is trained to predict the drug signatures of the target cell line given drug signatures from the reference cell lines. Some examples include D-GEX (Chen et al., 2016), GGAN (Wang et al., 2018), GeneDNN, GeneGAN, GeneLASSO, and SampleLASSO (Mancuso et al., 2020). D-GEX employs a multi-task multi-layer feedforward neural network model, whereas GGAN uses a conditional generative adversarial network. GeneDNN and GeneGAN address a gene-wise data imputation, where deep neural network and generative adversarial network model, respectively, are built for imputing the missing expression data of some genes. Finally, GeneLASSO and SampleLASSO are two related methods that rely on LASSO regression modeling (Tibshirani, 1996) to impute missing expression data for genes and samples, respectively. Since the above methods are based on supervised learning, they require training data that comprise drug signatures of the reference and target cell lines from the same set of conditions.

With the exception of TT-WOPT, the drug signature prediction considered in this work cannot be addressed by the imputation methods mentioned above since there is no overlapping drug treatment conditions between the reference and background transcriptomic data. Besides, when for the case of missing or unseen cells, the method TT-WOPT produced predictions with poor accuracy scores (Iwata et al., 2019). To address the lack of available algorithms, we developed CrossTx (Cross-cell line Transcriptomic signature prediction) for predicting transcriptomic drug signatures of an unseen target cell line, given drug signatures from a set of reference cell lines and background gene expression data of the target cells. Considering the features of the CMap dataset, we considered drug load as a covariate in generating drug signature predictions. CrossTx involves two steps: Predictor and Corrector. Given a drug of interest at a specific drug load, the Predictor produces cell line-agnostic drug signatures by averaging or a linear regression model using the reference drug signatures. The Corrector is based on the idea of correcting cell line-agnostic signatures by projection onto the transcriptomic latent space of the target cell line to obtain target-specific drug signatures. The transcriptomic latent space is constructed from the background (unlabeled) gene expression data of the target cell line using Principal Component Analysis (PCA) and/or Autoencoder (AE). For demonstration, we applied different Predictor-Corrector combinations to the CMap dataset using a Leave-One-Cell-line-Out (LOCO) procedure and assessed the accuracies of the predicted drug signatures using the Pearson correlation coefficient (*ρ*) and the area under Receiver Operating Characteristics and Precision-Recall curve, denoted by AUROC and AUPR, respectively.

## 2. Methods and Materials

### 2.1 Prediction of Drug Signature by CrossTx

CrossTx comprises two steps, a predictor and a corrector, that are applied in sequence. The Predictor (see Fig. 1b) generates a cell line-agnostic transcriptome signature for a drug at a specific drug load—defined as the drug concentration multiplied by the treatment duration—via averaging or linear regression model, hereon referred to as the ‘Mean’ and ‘Regression’ method, respectively. The Corrector subsequently projects the cell line-agnostic signature to the transcriptomic latent space of the target cell line via PCA and/or AE method to produce target-specific signature (see **Fig. 1c**). The details of the Predictor and Corrector steps are provided below.

Given a drug and drug load, the Predictor produces a vector of cell line-agnostic gene expression 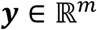 where *m* is the number of genes. The Mean method produces a cell line-agnostic gene expression value ***y*** = ***μ**_d,l_* by averaging the gene expression vectors in the reference dataset for the specified drug *d* and drug load *l* (see **Fig. 1b**). An obvious drawback of the Mean method is that it is unable to produce any prediction when the reference dataset does not have samples for the specified drug and drug load. In such a case, the Regression method should be used.

The Regression method relies on a linear regression model *r_g,d_*(*l*), in which the expression of gene *g* is the dependent variable and the drug load *l* is the independent variable, according to the following formulation:

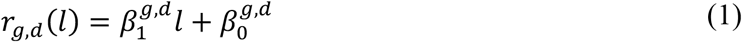

where 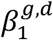 and 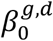 are the slope and intercept. Note that a separate linear regression model is built for each gene *g* and drug *d*. The final outcome is put into a vector of gene expression ***y***. In the application to the CMap dataset, the intercept was set to zero since the drug signatures have been normalized (z-scores) with a zero mean. The unknown slope was obtained using ordinary least squares method (Beck et al., 1977).

The first Corrector uses PCA as a projection method. The transcriptomic latent space of the target cell line is anticipated to be highly complex, limiting the ability of PCA to represent the complete latent space well. As illustrated in **Fig. 1c**, only a subset of the background gene expression data of the target cell line—samples nearest to the cell line-agnostic expression from the Predictor ***y***—is used in the PCA projection. Specifically, a subset of *k* background transcriptional profiles with the highest correlation with ***y*** are chosen for the projection (default *k* = 5). PCA is applied to the selected background expression profiles, and the top *p* principal components (PCs) are finally used for projection. The parameter *p* is set to the minimum number of PCs that cumulatively explain at least a given percentage of the total variance (default 80%). The PCA projection onto the latent space is done as follows:

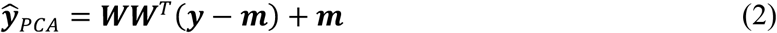

where 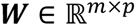 denotes the loading matrix for the selected PCs, 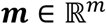 denotes the average expression of the selected background samples, and 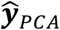 denotes the target-specific drug signature via PCA projection.

The second Corrector using AE employs the entire background data to learn transcriptomic latent representation of the target cell line. Autoencoders (AEs) and its many variants have been successfully applied to transcriptomic data for various purposes, from dimensionality reduction to imputation of missing values (Arisdakessian et al., 2019; Ding et al., 2018; Talwar et al., 2018; Wang & Gu, 2018). The AE architecture comprises an encoder and a decoder (see **Fig. 1c**), each is a perceptron with one hidden layer using the same number of hidden nodes. In CrossTx, the AE employs the hyperbolic tangent (tanh) activation function for both the encoder and the decoder. The number of nodes in the hidden layer *n_nodes_*, the dimension of the latent representation *n_z_*, the batch size *n_batch_* (*i.e*., the size of data subgrouping during the training), and the number of epochs *n_epoch_* (*i.e*., the number of iterations during training) are hyperparameters that need to be optimized. The encoder and the decoder are trained together to minimize the Mean Squared Error Φ:

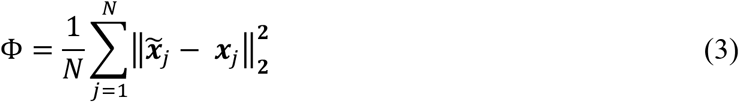

Here, 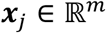 denotes a vector of background gene expression, 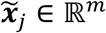 denotes the reconstructed gene expression by the AE, *N* is the number of the samples, and ||·||_2_ denotes the L-2 norm. Once trained, the output of the Predictor ***y*** is passed through the AE, as illustrated in **Fig. 1c**, to produce the target-specific gene expression 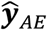.

### 2.2. Application to the CMap Dataset

The CMap drug transcriptomic signature dataset comprises gene expression data for 978 landmark genes measured using the L1000 assay (Subramanian et al., 2017). For assessing the performance of the proposed CrossTx method, we took the processed CMap drug signatures from a previous study by Pham et al. (Pham et al., 2021). This dataset was generated using a different peak deconvolution procedure from the original CMap, specifically using a Bayesian-based approach that generates more robust expression profiles (Qiu et al., 2020). The dataset includes seven cell lines with the most samples in the CMap, namely MCF7, A375, HT29, PC3, HA1E, YAPC, and HELA, and samples from six most frequent drug concentrations taken after 24 hours of treatment. The sample sizes for each cell line are given in **Table 1**.

**Table 1.** Sample Size of CMap Dataset in Performance Assessment. The number of test samples for the Mean is different from that for the Regression method since the Mean method requires at least one sample in the reference data for a given drug load.

| Sample size | MCF7 | A375 | HT29 | PC3 | HA1E | YAPC | HELA |
| --- | --- | --- | --- | --- | --- | --- | --- |
| Total | 1000 | 817 | 596 | 833 | 792 | 456 | 800 |
| Test: Mean / Regression | 250 / 253 | 354 / 377 | 263 / 272 | 240 / 246 | 267 / 280 | 209 / 210 | 173 / 174 |

For training the AE-based corrector, we performed hyperparameter tuning using Bayesian optimization method (Snoek et al., 2012) and identified the following optimal settings: *n_nodes_* of 150, *n_z_* of 100, *n_batch_* of 40, and *n_epoch_* of 300. We trained the AE using Adam optimizer (Kingma & Ba, 2014) with a learning rate of 10^−3^. We implemented the AE using Keras (version 2.10.0) (Chollet, 2015) with a TensorFlow backend (version 2.10.0) (Abadi, 2016).

For performance assessment, we followed Leave-One-Cell line-Out (LOCO) procedure. Briefly, we picked one of the cell lines in the dataset as the target cell line and the remaining six as the reference cell lines. Correspondingly, we treated the data from the assigned target cell line as the background transcriptome, and the remainder of the data as the reference drug signatures. We repeated the above procedure for each cell line in the dataset. Given a target cell line, we assessed the drug signature prediction accuracy (see Performance Scoring section) for the top 100 drugs in the dataset. The test sample size of the drug signature predictions for each target cell line is provided in **Table 1**. Since the Mean method is unable to produce cell line-agnostic signature when the reference dataset does not have any sample from the drug and drug load of interest, the test sample size for the Mean method was smaller than those for the Regression. But, the differences in the test sample sizes were small (average of 2.7%)—the largest difference is 6.1% for cell line A375. Note that when predicting the gene expression signature for a drug treatment, we excluded all samples of the respective drug from the background data—that is, the background data of the target cell line did not include any samples from the query drug.

### 2.3. Performance Scoring

The first accuracy score is the Pearson correlation coefficient *ρ*. Given the ground truth drug signature 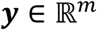 and the predicted signature 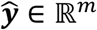, *ρ* is calculated as follows:

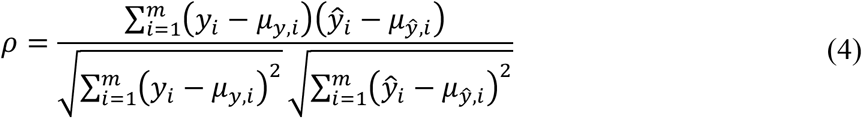

where ***μ**_y_* and 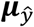 denote the average of the elements of ***y*** and 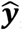, respectively (*i.e*., 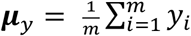). A higher *ρ* value suggests a more accurate prediction.

The next set of accuracy scores are the area under the Receiver Operating Characteristics (ROC)—the curve of true positive rate (TPR) vs. false positive rate (FPR)—(AUROC) and the area under Precision-Recall (AUPR) curve. These scores are computed for genes that up- or downregulated by drugs separately. For computing the AUROC and AUPR of upregulated (downregulated) genes, a ranked list of genes is created by sorting the genes based on the predicted gene expression in decreasing (increasing) order, from the most positive (negative) to the most negative (positive). The ROC and PR curves are generated by computing the numbers of true positive (TP), false positive (FP), false negative (FN), and true negative (TN) among the top *k* genes in the ranked list, for increasing *k*. TPs correspond to the intersection of the top *k* genes and the ground truth gene set (up-downregulated genes), while FPs are those among the top *k* genes that are not in the ground truth. Genes that are not in the top *k* of the list but are in the ground truth gene set, are the set of FNs. Finally, genes that are not among the top *k* nor in the ground truth are TNs. For ROC, the TPR and FPR are computed as follows:

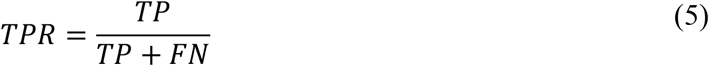

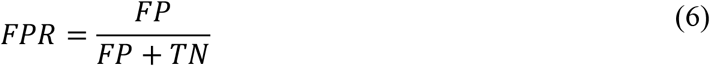

The ROC curve is constructed by plotting TPR versus FPR for increasing *k*. Meanwhile, for the PR curve, the precision and the recall are computed according to:

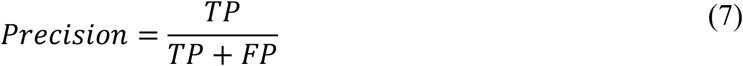

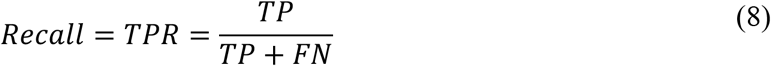

As the name suggests, the PR curve is the plot of Precision versus Recall for increasing *k*.

AUROC and AUPR values range between 0 and 1, where a value of 1 corresponds to a perfect prediction. Note that a random (binary) predictor has an expected value of AUROC equal to 0.5. The expected value of AUPR for a random prediction is equal to the proportion of positives (*i.e*., the number of genes in the ground truth over the total number of genes *m*). When the number genes in the ground truth set is much smaller than the total genes—i.e., when classes are highly imbalanced—the AUPR is a more informative metric of accuracy than the AUROC as it accounts for the ratio between positives and negatives (Saito & Rehmsmeier, 2015).

## 3. Results

CrossTx is a method for predicting drug transcriptional signatures of an unseen target cell line given data from reference cell lines and background gene expression data of the target cells. Background data of the target cells are considered unlabeled—that is, the conditions used in generating the data are not required nor used for drug signature predictions. The drug signature prediction of interest is unlike the common imputation since the gene expression data of the reference cell lines and the background data of the target cell lines, highlighted in blue and green in **Fig. 1a**, respectively, do not have any common conditions. The method CrossTx comprises two steps: a Predictor and a Corrector. The Predictor uses the reference dataset to produce cell line-agnostic transcriptional signatures via averaging or by linear regression (see **Fig. 1b**). The Corrector leverages the background dataset to capture the transcriptomic latent space of the target cell line using either PCA and/or an autoencoder (see **Fig. 1c**). In essence, the Corrector projects cell line-agnostic signatures from the Predictor onto this latent space to produce target-specific signatures. In the following, we assessed the performance of CrossTx and compared the method with TT-WOPT (Iwata et al., 2019). As noted earlier, the majority of existing imputation methods cannot be applied to the problem described in **Fig. 1a**.

We evaluated the accuracy of CrossTx predictions for drug signatures in seven cancer cell lines: MCF7, A375, HT29, PC3, HA1E, YAPC, and HELA. More specifically, we obtained preprocessed and filtered drug signatures from the study by Pham et al. (Pham et al., 2021). For testing, we followed the LOCO procedure (see Methods) where we systematically picked one cell line as the target and treated the remaining six cell lines as the reference. We generated drug signature predictions for 100 drugs with the most samples in the dataset (see **Table 1**). When making a signature prediction for a drug, we removed all samples of this drug from the background data of the target cells. For performance scoring, we employed the Pearson correlation *ρ* and the area under ROC (AUROC) and PR (AUPR) curve. When using the AUROC and AUPR, we performed the accuracy assessment for predictions of up- and down-regulated genes separately. In total, ten different drug signatures generated by the CrossTx were assessed: two cell line-agnostic signatures using either the Mean or the Regression method, and eight different combinations of the Predictor-Corrector: (Mean or Regression) + (PCA or AE or PCA + AE or AE + PCA).

**Fig. 2** gives the performance scores of CrossTx and TT-WOPT for all drug signature predictions across all target cell lines in LOCO procedure. The scores for individual target cell lines are given in **Table 2** (see **Supplementary Tables S1-5** for full results). The results show that cell line-agnostic drug signatures produced by the Mean and Regression methods have reasonable agreement with the ground truth data with correlations *ρ* averaging at 0.59 / 0.55 (Mean / Regression), AUROCs at 0.79 / 0.77 for up- and 0.80 /0.78 for downregulated genes, and AUPRs at 0.65 / 0.63 for up- and 0.67 / 0.65 for downregulated genes. In general, the Mean method gives a better accuracy than the Regression method (*p*-values <10^−4^, two-sided paired t-test). Still, the Regression method has the advantage over the Mean method in that it can provide predictions for drug load values that are not in the reference dataset. The Corrector using PCA projection improved the cell line-agnostic drug signatures from the Mean (Mean + PCA) and the Regression method (Regression + PCA). In contrast, the Corrector using AE provided accuracy improvements only for cell line-agnostic signatures from the Mean method (Mean + AE). When starting with the relatively poorer cell line-agnostic signatures from the Regression method, the AE projection (Regression + AE) actually degraded the accuracy of the drug signature predictions.

**Fig. 2.**
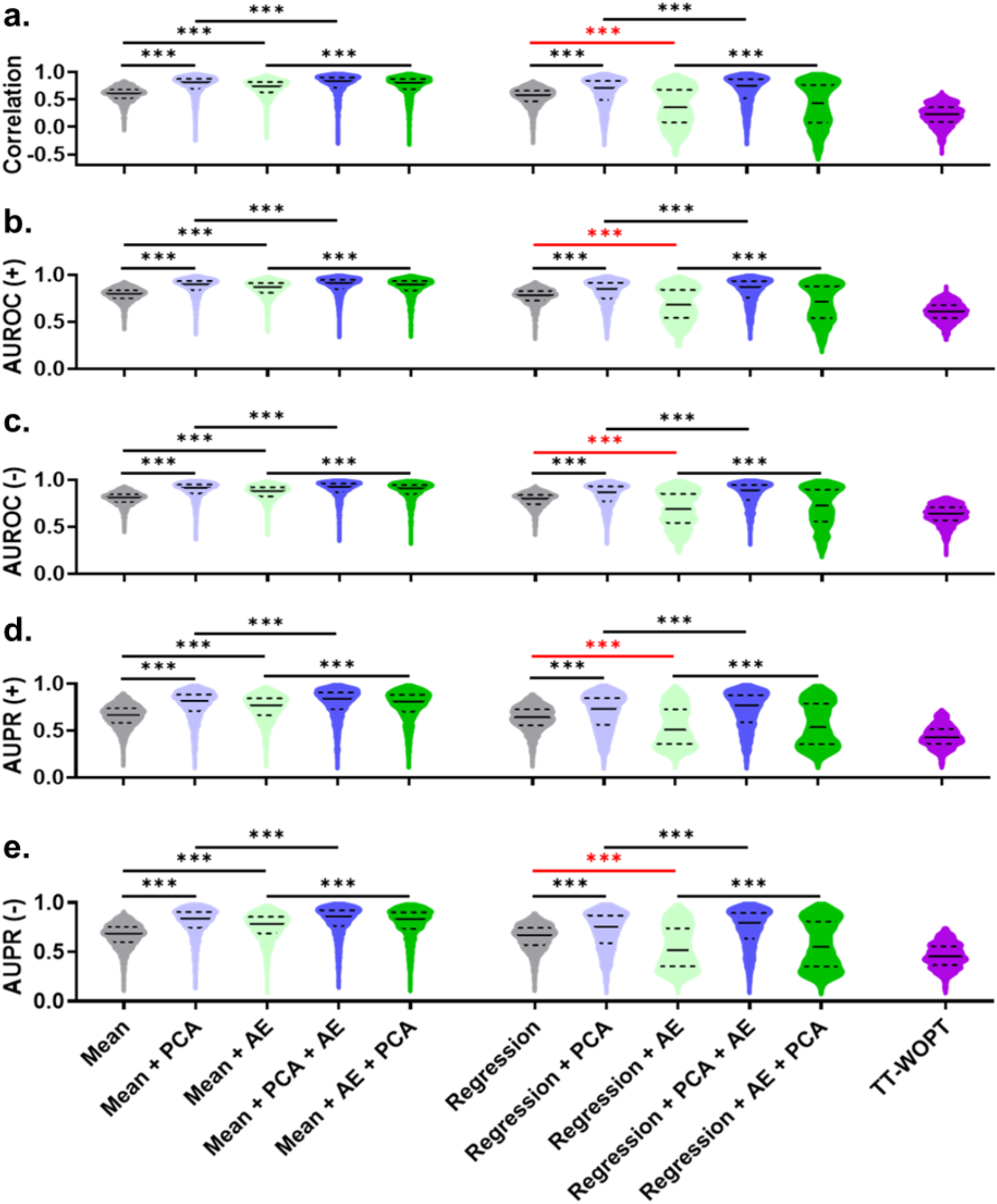
Accuracy of drug signature predictions by CrossTx and TT-WOPT. The accuracy of drug signature predictions was assessed by (**a**) Pearson correlation *ρ*, (**b-c**) AUROC, and (**d-e**) AUPR. AUROC (+) / AUROC (-) refer to values for up- / downregulated genes, respectively. Similarly, AUPR (+) / AUPR (-) refer to values for up- / downregulated genes, respectively. Solid lines correspond to medians and dashed lines give the inter-quartile range. Statistical significance was performed using two-sided paired t-test. Black *** indicates p-values < 10^−3^ for improved accuracy. Red *** indicates p-values < 10^−3^ for decreased accuracy.

**Table 2.**
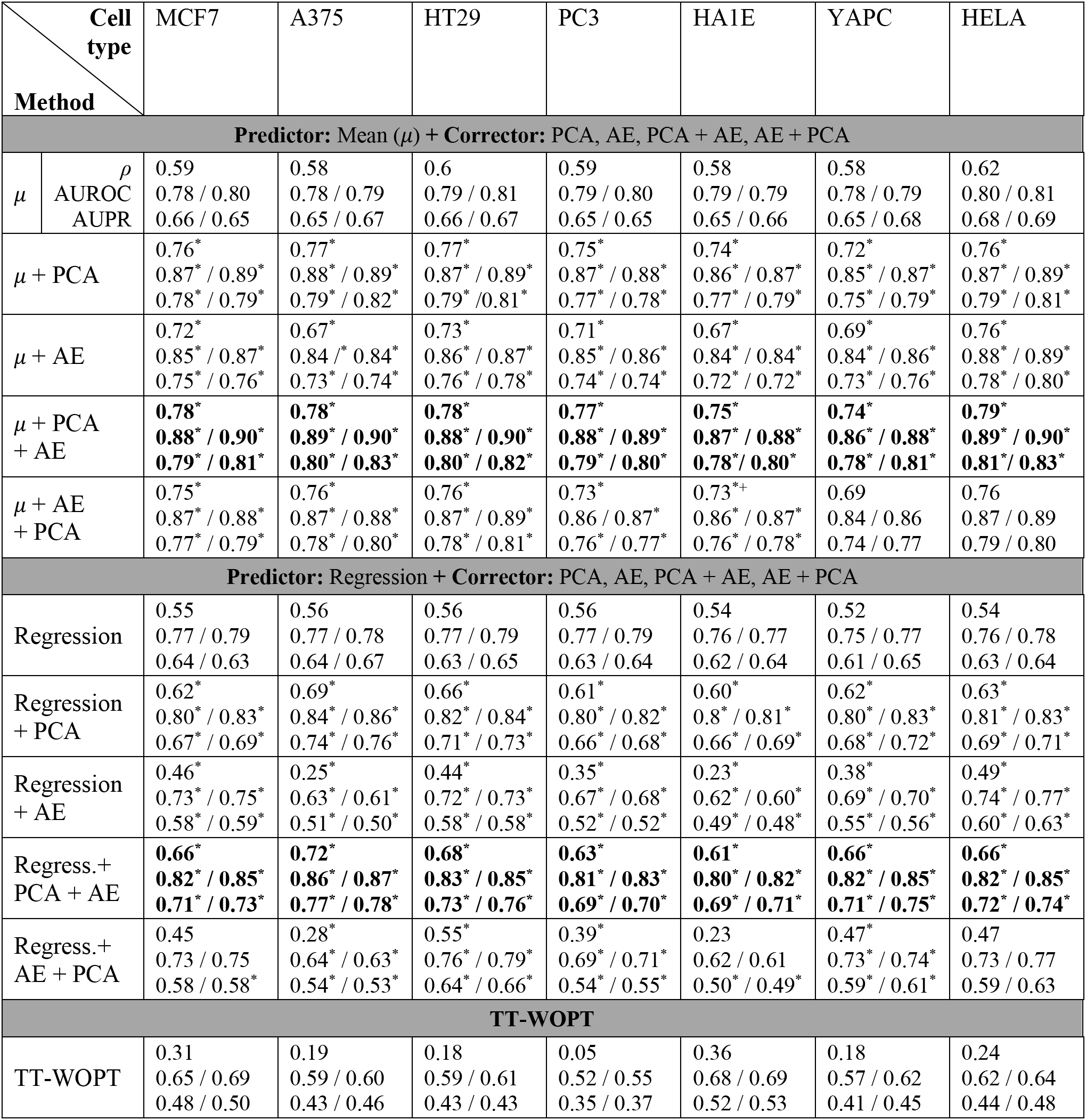
Accuracy of drug signature predictions by CrossTx and TT-WOPT for individual cell lines. For each method, the accuracy of drug signature predictions is assessed using Pearson correlation *ρ* (first row), AUROC (second row), and AUPR (third row). The two values of AUROCs correspond to the accuracy of predicting up- / downregulated genes. Similarly, the two values of AUROCs correspond to the accuracy of predicting up- / downregulated genes. Bold values highlight the best performing method for the respective accuracy metric. Statistical significance was established by two-sided paired t-test to assess the change in accuracy by adding a Corrector. For example, Mean + PCA was compared to Mean, while Mean + PCA + AE was compared to Mean + PCA. Note that the addition of a Corrector may degrade accuracy when using the AE method. *: *p*-values < 0.05.

The observed trend for the AE suggests that the AE projection method is a viable Corrector strategy only when using cell line-agnostic signatures that do not deviate far from the background transcriptomic latent space. This observation motivates combining the two Corrector methods in series. We tested whether the accuracy of the PCA projected drug signatures can be improved further by the AE, a strategy labelled as PCA + AE. The basic premise here is that the PCA projection would have brought the cell line-agnostic prediction close to the transcriptomic latent space of the target cells, for which the AE is suitable and may offer further improvements. Indeed, in every target cell line considered (see **Table 2**) and regardless of the method used to generate the cell line-agnostic signatures, the PCA + AE combination gave significant accuracy improvements over only using the PCA. For the sake of completeness, we also tested the AE + PCA combination as the Corrector—passing cell line-agnostic signatures to the AE, and then applying the PCA projection to the output of the AE. In this case, the PCA projection did not always provide improvements to the drug signatures produced by the AE. Overall, the best performing method was the Mean + PCA + AE combination. Lastly, the accuracy of TT-WOPT generated drug predictions were consistently poorer than the CrossTx predictions, including the drug signatures using the Mean method (*p*-values <10^−4^, two-sided paired t-test).

## 4. Discussion

Our work is motivated by the practical problem of predicting drug signature data in disease-relevant cells. Specifically, we addressed the challenge of leveraging large drug signature datasets, such as the CMap, to predict drug response in unseen cell lines—cells that are in the dataset. Since distinct gene, signaling, and metabolic network operate in different tissues and cell lines (Marbach et al., 2016; Schultz & Qutub, 2016; Sharma & Petsalaki, 2019), one expects that the molecular signatures of a drug will exhibit cell context specificity. For applications in drug discovery, drug repurposing, and precision medicine, there is a clear need for methods that are able to predict cell line-specific drug response accurately. We developed CrossTx to address the gap in the available algorithms for predicting transcriptional signatures of unseen cells. The method comprises two key components: a Predictor that generates cell line-agnostic drug signatures using a reference dataset and a Corrector that produces target-specific signatures by projecting cell line-agnostic signatures onto the transcriptomic latent space of a target cell line. Here, we tested two alternative methods for the Predictor (Mean and Regression) and two different latent projections for the Corrector (PCA and AE). But, this strategy is fully generalizable to other algorithms for the Predictor and the Corrector. For example, nonlinear regression models can be employed as the Predictor for generating cell line-agnostic gene expression signatures, while other machine learning approach for reconstructing and projection onto latent space can be adopted as the Corrector.

We tested the accuracy of ten different predictions that are produced by CrossTx using various combinations of Predictor-Corrector algorithms. We applied CrossTx to drug signatures from seven cell lines with the most samples in the CMap dataset. We found that averaging reference drug signatures (*i.e*., the Mean method) is a simple and efficacious method for the Predictor. When no reference samples at the specified drug load exist, the Regression method is a viable alternative method for a Predictor. Between the two Corrector methods, our tests showed that the PCA is superior to the AE, not only in terms of the accuracy of the resulting drug signatures, but also in its computational simplicity and robustness with respect to the accuracy of its input. As a Corrector, the AE projection gave improvements when the input signatures were similar to the background data of the target cell line. This observation is not surprising considering that the AE was trained using transcriptional data of the target cells. The combination of PCA and AE where PCA-projected drug signatures were inputted to the AE, gave the highest accuracy. But, the reverse implementation—applying AE projection and then PCA—did not outperform using only the PCA projection. Finally, all drug signatures generated by CrossTx had higher accuracies than those produced by an existing method called TT-WOPT that is based on tensor decomposition.

## 5. Conclusion

In summary, we presented an efficacious and generalizable predictor-corrector strategy for leveraging large reference molecular profiles of drug treatments (such as the CMap) to predict drug signatures in an unseen cell line—cells that are not included in the reference dataset. Our method CrossTx addresses a practical imputation problem where background transcriptional data of a target cell line exist, but these data were collected in experiments that did not have any overlap with those from the reference data. Of note, CrossTx does not require nor utilize any information related to the experimental conditions of the background data. To the best of our knowledge, none of the existing imputation algorithms, specifically those developed for the CMap dataset, are applicable to the problem at hand (see **Fig. 1a**). While a comparison to the method TT-WOPT was performed, TT-WOPT was not able to take advantage background data of the target cell line (*i.e*., the method was not developed nor optimized for our problem). Lastly, despite the focus on drug signatures, the prediction problem and the proposed predictor-corrector strategy are applicable to many practical scenarios where one would like to estimate the transcriptional response of a cell line to a treatment/perturbation, given gene expression data of other cell lines exposed to the same treatment and data of the cell line of interest from the literature or other own experiments.

## Acknowledgments

This work was supported by start-up funding from University at Buffalo-SUNY and funding from NSF HDR I-DIRSE-Ideas Labs (grant #1940162).

