## Supplementary Tables for "CrossTx: Cross-cell line Transcriptomic Signature Predictions"

**Supplementary Table S1. Pearson correlations of drug signature predictions by CrossTx and TT-WOPT for individual cell lines.** Values correspond to mean ± standard deviation. Bold values signify the best method based on Pearson correlation. Statistical significance was established by two-sided paired t-test to assessed the change in accuracy by adding a Corrector. For example, Mean + PCA was compared to Mean, while Mean + PCA + AE was compared to Mean + PCA. Note that the addition of a Corrector may degrade accuracy. TT-WOPT was compared to the Mean method. ∗: $p$-values < 0.05.

| Method \ Cell type | MCF7 | A375 | HT29 | PC3 | HA1E | YAPC | HELA |
|---|---|---|---|---|---|---|---|
| **Predictor:** Mean ($\mu$) **+ Corrector:** PCA, AE, PCA + AE, AE + PCA | | | | | | | |
| Mean | 0.59 ±0.14 | 0.58 ±0.12 | 0.6 ±0.13 | 0.59 ±0.14 | 0.58 ±0.14 | 0.58 ±0.18 | 0.62 ±0.11 |
| Mean + PCA | 0.76* ±0.17 | 0.77* ±0.15 | 0.77* ±0.18 | 0.75* ±0.19 | 0.74* ±0.21 | 0.72* ±0.23 | 0.76* ±0.2 |
| Mean + AE | 0.72* ±0.15 | 0.67* ±0.15 | 0.73* ±0.14 | 0.71* ±0.16 | 0.67* ±0.16 | 0.69* ±0.22 | 0.76* ±0.12 |
| Mean + PCA + AE | **0.78* ±0.17** | **0.78* ±0.16** | **0.78* ±0.18** | **0.77* ±0.19** | **0.75* ±0.22** | **0.74* ±0.24** | **0.79* ±0.19** |
| Mean + AE + PCA | 0.75* ±0.18 | 0.76* ±0.16 | 0.76* ±0.18 | 0.73* ±0.21 | 0.73* ±0.22 | 0.69 ±0.25 | 0.76 ±0.2 |
| **Predictor:** Regression **+ Corrector:** PCA, AE, PCA + AE, AE + PCA | | | | | | | |
| Regression | 0.55 ±0.15 | 0.56 ±0.14 | 0.56 ±0.17 | 0.56 ±0.16 | 0.54 ±0.16 | 0.52 ±0.22 | 0.54 ±0.16 |
| Regression + PCA | 0.62* ±0.25 | 0.69* ±0.22 | 0.66* ±0.25 | 0.61* ±0.26 | 0.6* ±0.27 | 0.62* ±0.27 | 0.63* ±0.27 |
| Regression + AE | 0.46* ±0.26 | 0.25* ±0.38 | 0.44* ±0.3 | 0.35* ±0.34 | 0.23* ±0.41 | 0.38* ±0.31 | 0.49* ±0.29 |
| Regression + PCA + AE | **0.66* ±0.24** | **0.72* ±0.22** | **0.68* ±0.26** | **0.63* ±0.26** | **0.61* ±0.29** | **0.66* ±0.26** | **0.66* ±0.26** |
| Regression + AE + PCA | 0.45 ±0.3 | 0.28* ±0.47 | 0.55* ±0.31 | 0.39* ±0.35 | 0.23 ±0.46 | 0.47* ±0.29 | 0.47 ±0.31 |
| **TT-WOPT** | | | | | | | |
| TT-WOPT | 0.31* ±0.16 | 0.19* ±0.2 | 0.18* ±0.18 | 0.05* ±0.12 | 0.36* ±0.15 | 0.18* ±0.15 | 0.24* ±0.2 |

**Supplementary Table S2. AUPRs of drug signature predictions by CrossTx using the Mean method as Predictor and TT-WOPT for individual cell lines.** Values correspond to mean ± standard deviation. Bold values signify the best method based on Pearson correlation. Statistical significance was established by two-sided paired t-test to assessed the change in accuracy by adding a Corrector. For example, Mean + PCA was compared to Mean, while Mean + PCA + AE was compared to Mean + PCA. Note that the addition of a Corrector may degrade accuracy. TT-WOPT was compared to the Mean method. ∗: $p$-values < 0.05.

| **Method** \ **Cell type** | MCF7 | A375 | HT29 | PC3 | HA1E | YAPC | HELA |
|---|---|---|---|---|---|---|---|
| **AUPR for predicting upregulated genes** | | | | | | | |
| Mean | 0.66 ±0.11 | 0.65 ±0.1 | 0.66 ±0.11 | 0.65 ±0.12 | 0.65 ±0.12 | 0.65 ±0.14 | 0.68 ±0.11 |
| Mean + PCA | 0.78* ±0.14 | 0.79* ±0.13 | 0.79* ±0.15 | 0.77* ±0.15 | 0.77* ±0.17 | 0.75* ±0.17 | 0.79* ±0.16 |
| Mean + AE | 0.75* ±0.13 | 0.73* ±0.13 | 0.76* ±0.14 | 0.74* ±0.14 | 0.72* ±0.14 | 0.73* ±0.17 | 0.78* ±0.12 |
| Mean + PCA + AE | **0.79* ±0.14** | **0.80* ±0.13** | **0.80* ±0.15** | **0.79* ±0.16** | **0.78* ±0.17** | **0.78* ±0.17** | **0.81* ±0.15** |
| Mean + AE + PCA | 0.77* ±0.14 | 0.78* ±0.13 | 0.78* ±0.15 | 0.76* ±0.17 | 0.76* ±0.17 | 0.74 ±0.18 | 0.79 ±0.15 |
| TT-WOPT | 0.48* ±0.1 | 0.431* ±0.095 | 0.428* ±0.095 | 0.347* ±0.087 | 0.52* ±0.12 | 0.405* ±0.097 | 0.44* ±0.13 |
| **AUPR for predicting downregulated genes** | | | | | | | |
| Mean | 0.65 ±0.11 | 0.67 ±0.11 | 0.67 ±0.11 | 0.65 ±0.13 | 0.66 ±0.13 | 0.68 ±0.12 | 0.686 ±0.096 |
| Mean + PCA | 0.79* ±0.14 | 0.82* ±0.13 | 0.81* ±0.14 | 0.78* ±0.16 | 0.79* ±0.17 | 0.79* ±0.15 | 0.81* ±0.15 |
| Mean + AE | 0.76* ±0.12 | 0.74* ±0.13 | 0.78* ±0.13 | 0.74* ±0.15 | 0.72* ±0.14 | 0.76* ±0.15 | 0.80* ±0.1 |
| Mean + PCA + AE | **0.81* ±0.14** | **0.83* ±0.13** | **0.82* ±0.14** | **0.80* ±0.16** | **0.80* ±0.17** | **0.81* ±0.16** | **0.83* ±0.14** |
| Mean + AE + PCA | 0.79* ±0.15 | 0.80* ±0.13 | 0.81* ±0.14 | 0.77* ±0.17 | 0.78* ±0.17 | 0.77 ±0.16 | 0.80 ±0.15 |
| TT-WOPT | 0.50* ±0.11 | 0.46* ±0.12 | 0.43* ±0.11 | 0.365* ±0.097 | 0.53* ±0.14 | 0.448* ±0.087 | 0.48* ±0.12 |

**Supplementary Table S3. AUPRs of drug signature predictions by CrossTx using the Regression method as Predictor and TT-WOPT for individual cell lines.** Values correspond to mean ± standard deviation. Bold values signify the best method based on Pearson correlation. Statistical significance was established by two-sided paired t-test to assessed the change in accuracy by adding a Corrector. For example, Mean + PCA was compared to Mean, while Mean + PCA + AE was compared to Mean + PCA. Note that the addition of a Corrector may degrade accuracy. TT-WOPT was compared to the Regression method. ∗: $p$-values < 0.05.

| **Cell type** / **Method** | MCF7 | A375 | HT29 | PC3 | HA1E | YAPC | HELA |
|---|---|---|---|---|---|---|---|
| **AUPR for predicting upregulated genes** | | | | | | | |
| Regression | 0.64 ±0.12 | 0.64 ±0.11 | 0.63 ±0.13 | 0.63 ±0.13 | 0.62 ±0.14 | 0.61 ±0.15 | 0.63 ±0.13 |
| Regression + PCA | 0.67* ±0.18 | 0.74* ±0.16 | 0.71* ±0.19 | 0.66* ±0.2 | 0.66* ±0.19 | 0.68* ±0.2 | 0.69* ±0.19 |
| Regression + AE | 0.58* ±0.18 | 0.51* ±0.21 | 0.58* ±0.2 | 0.52* ±0.22 | 0.49* ±0.22 | 0.55* ±0.2 | 0.60* ±0.2 |
| Regression + PCA + AE | **0.71*** **±0.18** | **0.77*** **±0.17** | **0.73*** **±0.2** | **0.69*** **±0.2** | **0.69*** **±0.2** | **0.71*** **±0.2** | **0.72*** **±0.19** |
| Regression + AE + PCA | 0.58 ±0.19 | 0.54* ±0.25 | 0.64* ±0.22 | 0.54* ±0.23 | 0.50* ±0.24 | 0.59* ±0.2 | 0.59 ±0.22 |
| TT-WOPT | 0.48* ±0.1 | 0.431* ±0.095 | 0.428* ±0.095 | 0.347* ±0.087 | 0.52* ±0.12 | 0.405* ±0.097 | 0.44* ±0.13 |
| **AUPR for predicting downregulated genes** | | | | | | | |
| Regression | 0.63 ±0.12 | 0.67 ±0.11 | 0.65 ±0.12 | 0.64 ±0.14 | 0.64 ±0.14 | 0.65 ±0.14 | 0.64 ±0.12 |
| Regression + PCA | 0.69* ±0.19 | 0.76* ±0.16 | 0.73* ±0.18 | 0.68* ±0.2 | 0.69* ±0.19 | 0.72* ±0.19 | 0.71* ±0.19 |
| Regression + AE | 0.59* ±0.18 | 0.5* ±0.22 | 0.58* ±0.21 | 0.52* ±0.22 | 0.48* ±0.22 | 0.56* ±0.21 | 0.63* ±0.19 |
| Regression + PCA + AE | **0.73*** **±0.18** | **0.78*** **±0.16** | **0.76*** **±0.19** | **0.70*** **±0.2** | **0.71*** **±0.2** | **0.75*** **±0.18** | **0.74*** **±0.18** |
| Regression + AE + PCA | 0.58* ±0.21 | 0.53* ±0.26 | 0.66* ±0.22 | 0.55* ±0.23 | 0.49* ±0.26 | 0.61* ±0.2 | 0.63 ±0.2 |
| TT-WOPT | 0.50* ±0.11 | 0.46* ±0.12 | 0.43* ±0.11 | 0.365 * ±0.097 | 0.53* ±0.14 | 0.448* ±0.087 | 0.48 * ±0.12 |

**Supplementary Table S4. AUROCs of drug signature predictions by CrossTx using the Mean method as Predictor and TT-WOPT for individual cell lines.** Values correspond to mean ± standard deviation. Bold values signify the best method based on Pearson correlation. Statistical significance was established by two-sided paired t-test to assessed the change in accuracy by adding a Corrector. For example, Mean + PCA was compared to Mean, while Mean + PCA + AE was compared to Mean + PCA. Note that the addition of a Corrector may degrade accuracy. TT-WOPT was compared to the Mean method. ∗: $p$-values < 0.05.

| Method \ Cell type | MCF7 | A375 | HT29 | PC3 | HA1E | YAPC | HELA |
|---|---|---|---|---|---|---|---|
| **AUROC for predicting upregulated genes** | | | | | | | |
| Mean | 0.782 ±0.069 | 0.779 ±0.064 | 0.787 ±0.068 | 0.791 ±0.067 | 0.785 ±0.073 | 0.781 ±0.092 | 0.799 ±0.064 |
| Mean + PCA | 0.87* ±0.085 | 0.878* ±0.079 | 0.874* ±0.096 | 0.869* ±0.096 | 0.86* ±0.11 | 0.85* ±0.12 | 0.87* ±0.1 |
| Mean + AE | 0.854* ±0.078 | 0.835* ±0.082 | 0.858* ±0.08 | 0.854* ±0.08 | 0.835* ±0.085 | 0.84* ±0.12 | 0.877* ±0.069 |
| Mean + PCA + AE | **0.88* ±0.087** | **0.886* ±0.081** | **0.883* ±0.098** | **0.881* ±0.097** | **0.87* ±0.11** | **0.86* ±0.12** | **0.89* ±0.1** |
| Mean + AE + PCA | 0.867* ±0.091 | 0.871* ±0.083 | 0.87* ±0.095 | 0.86 ±0.11 | 0.86* ±0.11 | 0.84 ±0.13 | 0.87 ±0.1 |
| TT-WOPT | 0.647* ±0.08 | 0.592* ±0.097 | 0.586* ±0.09 | 0.522* ±0.061 | 0.683* ±0.08 | 0.572* ±0.074 | 0.618* ±0.099 |
| **AUROC for predicting downregulated genes** | | | | | | | |
| Mean | 0.80 ±0.07 | 0.788 ±0.065 | 0.805 ±0.065 | 0.798 ±0.075 | 0.789 ±0.07 | 0.793 ±0.087 | 0.812 ±0.055 |
| Mean + PCA | 0.889* ±0.086 | 0.891* ±0.076 | 0.892* ±0.086 | 0.878* ±0.098 | 0.87* ±0.11 | 0.87* ±0.11 | 0.891* ±0.088 |
| Mean + AE | 0.872* ±0.078 | 0.841* ±0.081 | 0.874* ±0.075 | 0.857* ±0.089 | 0.838* ±0.085 | 0.86* ±0.11 | 0.892* ±0.059 |
| Mean + PCA + AE | **0.898* ±0.086** | **0.897* ±0.079** | **0.901* ±0.088** | **0.89* ±0.1** | **0.88* ±0.11** | **0.88* ±0.11** | **0.903* ±0.086** |
| Mean + AE + PCA | 0.884* ±0.092 | 0.883* ±0.081 | 0.888* ±0.087 | 0.87* ±0.11 | 0.87* ±0.11 | 0.86 ±0.12 | 0.889 ±0.09 |
| TT-WOPT | 0.686* ±0.089 | 0.60* ±0.11 | 0.605* ±0.096 | 0.545* ±0.064 | 0.694* ±0.076 | 0.621* ±0.075 | 0.643* ±0.098 |

**Supplementary Table S5. AUROCs of drug signature predictions by CrossTx using the Regression method as Predictor and TT-WOPT for individual cell lines.** Values correspond to mean ± standard deviation. Bold values signify the best method based on Pearson correlation. Statistical significance was established by two-sided paired t-test to assessed the change in accuracy by adding a Corrector. For example, Mean + PCA was compared to Mean, while Mean + PCA + AE was compared to Mean + PCA. Note that the addition of a Corrector may degrade accuracy. TT-WOPT was compared to the Regression method. ∗: *p*-values < 0.05.

| **Cell type** / **Method** | MCF7 | A375 | HT29 | PC3 | HA1E | YAPC | HELA |
|---|---|---|---|---|---|---|---|
| **AUROC for predicting upregulated genes** | | | | | | | |
| Regression | 0.765 ±0.078 | 0.774 ±0.07 | 0.767 ±0.089 | 0.772 ±0.079 | 0.764 ±0.086 | 0.75 ±0.11 | 0.764 ±0.088 |
| Regression + PCA | 0.80* ±0.12 | 0.84* ±0.11 | 0.82* ±0.13 | 0.80* ±0.13 | 0.80* ±0.13 | 0.80* ±0.14 | 0.81* ±0.14 |
| Regression + AE | 0.73* ±0.13 | 0.63* ±0.19 | 0.72* ±0.15 | 0.67* ±0.17 | 0.62* ±0.21 | 0.69* ±0.15 | 0.74* ±0.15 |
| Regression + PCA + AE | **0.82* ±0.12** | **0.86* ±0.11** | **0.83* ±0.14** | **0.81* ±0.13** | **0.8* ±0.15** | **0.82* ±0.14** | **0.82* ±0.14** |
| Regression + AE + PCA | 0.73 ±0.14 | 0.64* ±0.23 | 0.76* ±0.16 | 0.69* ±0.18 | 0.62 ±0.23 | 0.73* ±0.14 | 0.73 ±0.16 |
| TT-WOPT | 0.647 ±0.08 | 0.592 ±0.097 | 0.586 ±0.09 | 0.522 ±0.061 | 0.683 ±0.08 | 0.572 ±0.074 | 0.618 ±0.099 |
| **AUROC for predicting downregulated genes** | | | | | | | |
| Regression | 0.786 ±0.075 | 0.784 ±0.068 | 0.788 ±0.084 | 0.787 ±0.081 | 0.773 ±0.081 | 0.77 ±0.1 | 0.781 ±0.076 |
| Regression + PCA | 0.83* ±0.12 | 0.86* ±0.1 | 0.84* ±0.12 | 0.82* ±0.12 | 0.81* ±0.13 | 0.83* ±0.13 | 0.83* ±0.12 |
| Regression + AE | 0.75* ±0.13 | 0.61* ±0.2 | 0.73* ±0.15 | 0.68* ±0.18 | 0.60* ±0.22 | 0.70* ±0.16 | 0.77* ±0.13 |
| Regression + PCA + AE | **0.85* ±0.11** | **0.87* ±0.11** | **0.85* ±0.12** | **0.83* ±0.13** | **0.82* ±0.14** | **0.85* ±0.12** | **0.85* ±0.12** |
| Regression + AE + PCA | 0.75 ±0.15 | 0.63* ±0.25 | 0.79* ±0.15 | 0.71* ±0.17 | 0.61 ±0.24 | 0.74* ±0.15 | 0.77 ±0.14 |
| TT-WOPT | 0.686* ±0.089 | 0.60* ±0.11 | 0.605* ±0.096 | 0.545* ±0.064 | 0.694* ±0.076 | 0.621* ±0.075 | 0.643* ±0.098 |
